# Predicting absolute protein folding stability using generative models

**DOI:** 10.1101/2024.03.14.584940

**Authors:** Matteo Cagiada, Sergey Ovchinnikov, Kresten Lindorff-Larsen

## Abstract

While there has been substantial progress in our ability to predict changes in protein stability due to amino acid substitutions, progress has been slower in methods to predict the absolute stability of a protein. Here we show how a generative model for protein sequence can be leveraged to predict absolute protein stability. We benchmark our predictions across a broad set of proteins and find a mean error of 1.5 kcal/mol and a correlation coefficient of 0.7 for the absolute stability across a range of natural, small–medium sized proteins up to ca. 150 amino acid residues. We analyse current limitations and future directions including how such model may be useful for predicting conformational free energies. Our approach is simple to use and freely available via an online implementation.

## Introduction

The stability of a protein is a critical property that affects its biological functions, as well as our ability to engineer proteins for new purposes. This stability is mainly the result of a delicate balance between the atomic interactions in the folded state, the higher conformational entropy in the unfolded state, as well as a complex relationship between the protein structure and water properties (***Aninsen, 1973***; ***Fersht, 2017***). By protein stability, we here concern ourselves with the (thermodynamic) free energy of unfolding, Δ*G*, under a given set of conditions such as temperature and pH.

Experimentally, Δ*G* can be measured using a range of biophysical and spectroscopic techniques, though generally the value under native-like conditions requires extrapolations from for example high concentrations of denaturant or elevated temperatures (***Lindorff-Larsen and Teilum, 2021***). These measurements generally require purification of each variant and is relatively labour intensive, even though automated methods have been used (***Nisthal et al., 2019***). While the thermodynamic stability has been determined for a number of proteins and their variants using these approaches, the resulting data are relatively inhomogeneous in terms of methods used and experimental conditions (***Nikam et al., 2021***; ***Xavier et al., 2021***). As an alternative, protein stability may in certain cases also be indirectly quantified using kinetic experiments (***Park and Marqusee, 2005***), and this has recently been combined with large-scale DNA synthesis and display technologies to quantify protein stability at a very large scale (***Tsuboyama et al., 2023***). While this type of experiments has provided a wealth of information, it is for technical reasons currently limited to small proteins with less than about 80 amino acid residues and which fold in a two-state manner.

As a supplement to experimental measurements, computational predictions can be used to assess the stability of a protein. With few exceptions, these methods have mostly focused on predictions of *changes* in thermodynamic stability (ΔΔ*G*), typically between a protein and a variant that differs from it by one or very few amino acid substitutions. A number of methods to predict ΔΔ*G* have been developed (***Pancotti et al., 2022***), and their accuracy in part originates from a number of important compensating effects. These include that while Δ*G* may be sensitive to the conditions (pH, temperature, etc), ΔΔ*G* will be less sensitive. Similarly, while both Δ*G* and ΔΔ*G* contain contributions from many interactions across the protein, many of these will cancel out when comparing the stability of a wild-type protein and a variant with a single amino acid substitution. Thus, in these senses ΔΔ*G* is a more robust quantity than Δ*G*.

One additional complication in analysing and predicting protein stability is that many of the underlying thermodynamic parameters (Δ*H*, Δ*s* and Δ*C*_*p*_) are strongly correlated with the number of residues (length) of the protein (***Robertson and Murphy, 1997***). These effects in turn suggest that it will be easier to predict the thermal stability (melting temperature) of a protein than the stability at a fixed temperature (***Pucci and Rooman, 2014***), and indeed several methods have been developed to predict melting temperatures. Previous work used these properties to predict, for example, stability from chain length (***Ghosh and Dill, 2009***; ***De Sancho et al., 2009***) with more recent methods taking additional physical and statistical information into account (***Ruiz-Blanco et al., 2013***; ***Pucci and Rooman, 2014***; ***Yang et al., 2022***).

While progress in predicting stability (Δ*G*) has been relatively slow, developments in the field of machine learning have resulted in an increase in the number of prediction methods to estimate changes in protein stability (ΔΔ*G*) (***Pucci et al., 2022***). While these developments are encouraging, a recent example illustrates the difficulty of training models that learn both Δ*G* and ΔΔ*G* (***Chu and Siegel, 2023***). Specifically, the authors trained a model directly on large-scale measurements of Δ*G* for small proteins, and showed that the model could be used to estimate both the global stability of similar folds and the effect of substitutions. However, the work also showed limited transferability, suggesting that training broadly transferable models for Δ*G* on a set of small proteins may be difficult.

Our work here uses a generative model as input to predict protein stability. Generative models for proteins are deep learning architectures that can be used in protein design to generate novel protein sequences or structures. They are generally based on representations learned from self-supervised training on large sequence or structure databases. In this class of models, fixed-backbone design architectures use the backbone coordinates of a target fold to generate novel sequences that fit the given structure, implicitly maximizing the fitness of the amino acid side chains in the local protein environment. Among these models, the so-called ‘inverse folding’ approach of the Evolutionary Scale Model family (ESM-IF)((***Hsu et al., 2022***)) has been trained to recover and generate novel amino acid sequences given an input protein fold during training on millions of sequence-structure pairs. ESM-IF achieves this by using an invariant graph neural network followed by a generic transformer(***Vaswani, 2017***). First, the graph model uses rotation equivalence and invariant transformations to generate residue embeddings that represent the geometric features of a backbone for a target protein fold, and then the transformer module uses the generated embeddings to return amino acid likelihoods and sampled sequences. The model generates likelihoods and sequences autoregressively, decoding the amino acid from the N-to C-terminus and using the information from the previous sampled positions to condition the next predicted residue. It has been shown that the ESM-IF likelihoods can be used to assess ΔΔ*G* by comparing the likelihoods of the wild-type and variant sequence, using the wild-type structure as input (***Hsu et al., 2022***; ***Chen and Gong, 2022***; ***Reeves and Kalyaanamoorthy, 2023***; ***Notin et al., 2024***). Here, we explored whether models such as ESM-IF can also be used to solve the more difficult problem of predicting protein stability across different folds and sequences (Δ*G*). Our manuscript is structured as follows. First, we show how information from ESM-IF can be used to predict absolute stability for proteins with an equilibrium two-state folding mechanism. Second, we explore how well the model can assess the relative stability between two different configurations of a protein. Third, we show limitations for predicting stability for larger and more complex systems, where cooperativity between structural elements may also play a role. Finally, we briefly discuss issues that we believe need to be addressed by experiments and future models in order to advance the field.

## Results and Discussion

### Predicting global folding stability using generative models

It has been shown that the ESM-IF model can be used to predict changes in protein stability (***Hsu et al., 2022***; ***Chen and Gong, 2022***; ***Reeves and Kalyaanamoorthy, 2023***; ***Notin et al., 2024***), suggesting that it might encode some level of protein biophysics that relates protein structure to thermodynamics. We therefore hypothesized that the ESM-IF model could also be used to predict the absolute stability of a given fold (Δ*G*) given its sequence. Because protein stability may vary substantially across conditions, we created a benchmark data set of experimental Δ*G* measurements by merging a curated data set of Δ*G* measurements of two-state folding proteins obtained under ‘standard’ experimental conditions (25 °C, pH 7.0) (***Maxwell et al., 2005***) with Δ*G* measurements from a large-scale proteolysis experiment on small single-domain proteins (***Tsuboyama et al., 2023***) (performed at ca. 25 °C, pH 7.4). In total, the benchmark data include 265 proteins ranging in size from 30 to 150 residues (Fig. S1A) and Δ*G* ranging from 1–12 kcal/mol (Fig. S1B). The dataset includes (i) 26 proteins whose kinetics and thermodynamics have been studied biophysically and that range in size from 61–151 residues (***Maxwell et al., 2005***) and (ii) 239 naturally occurring wild-type proteins (40–72 residues long) from the recently described ‘megascale’ experiment (***Tsuboyama et al., 2023***).

We used this data set to assess whether conditional likelihoods for amino acids calculated from ESM-IF for a given structure could be used to predict its global stability. Since it has been shown that protein stability correlates with protein size (***Ghosh and Dill, 2009***; ***De Sancho et al., 2009***), we used as measure the sum of likelihoods for the target sequence 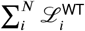 (Eq. 1). Here, the sum runs over the *N* residues in the protein, and 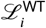 is the likelihood from ESM-IF for having the reference (wild-type) amino acid at position (see Methods). We calculated this measure for each of the 265 proteins and obtain a Pearson’s correlation of 0.69 with the experimental Δ*G* values, demonstrating how ESM-IF can be used to predict absolute protein stabilities (Fig. 1A). We then compared the accuracy achieved by Eq. 1 to other simpler measures such as the number of contacts in the fold or the sequence length (***Bastolla and Demetrius, 2005***; ***Ghosh and Dill, 2009***; ***De Sancho et al., 2009***). We find that both of these measures are substantially less correlated with the experimental Δ*G* values than Δ*G*_pred_ calculated from ESM-IF (Fig. 1B,C). We also assessed whether the summed model entropy could be used to predict protein stability and find that there was no correlation with experimental data (Fig. S2).

**Figure 1.**
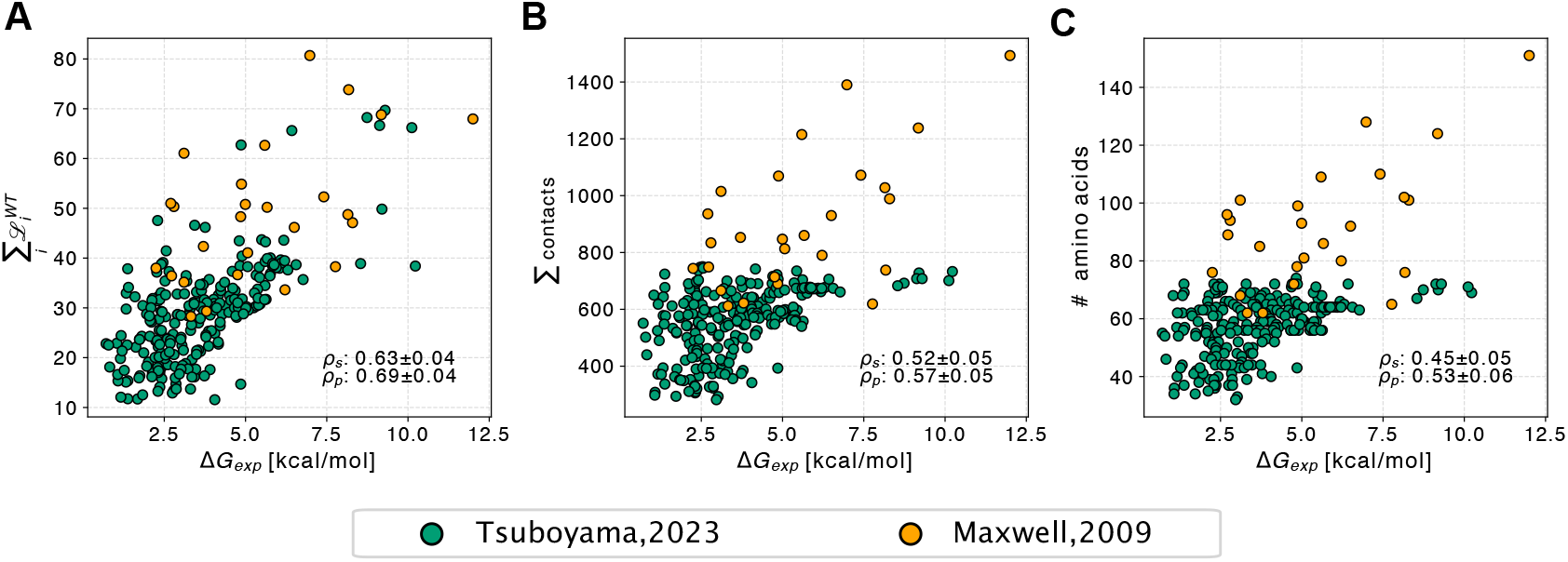
Predicting absolute protein stability. We show a comparison between Δ*G*_exp_ values for 265 proteins from two different data sets (represented by different colours) and (A) the sum of raw likelihoods predicted by ESM-IF, (B) the total number of contacts formed in the structure and, for comparison, (C) the number of residues. The Spearman’s and Pearson’s correlation coefficient are shown in each panel. The reported standard errors were obtained by bootstrapping.

We then convert the values from Eq. 1 into Δ*G*_pred_ in physical units, by fitting a scaling constant and offset using the experimental data (see Methods). The resulting predictions have a root-mean-square-error between Δ*G*_exp_ and Δ*G*_pred_ of 1.4 kcal/mol (Fig. 2), mostly constant across proteins of different stabilities (Fig. S3). This conversion from likelihoods to physical units involves fitting two parameters, and cross-validation shows the errors to be robust (Fig. S4), and lower than the baselines we examined (Table 1).

**Table 1.** RMSE comparison for the null model and different models, including ESM-IF, for the predictions fitted with experimental data on the experimental data, again.

| Method | Null model | Sequence model | ACN model | ESM-IF model |
| --- | --- | --- | --- | --- |
| RMSE | 1.88 kcal/mol | 1.55 kcal/mol | 1.53 kcal/mol | 1.39 kcal/mol |

**Figure 2.**
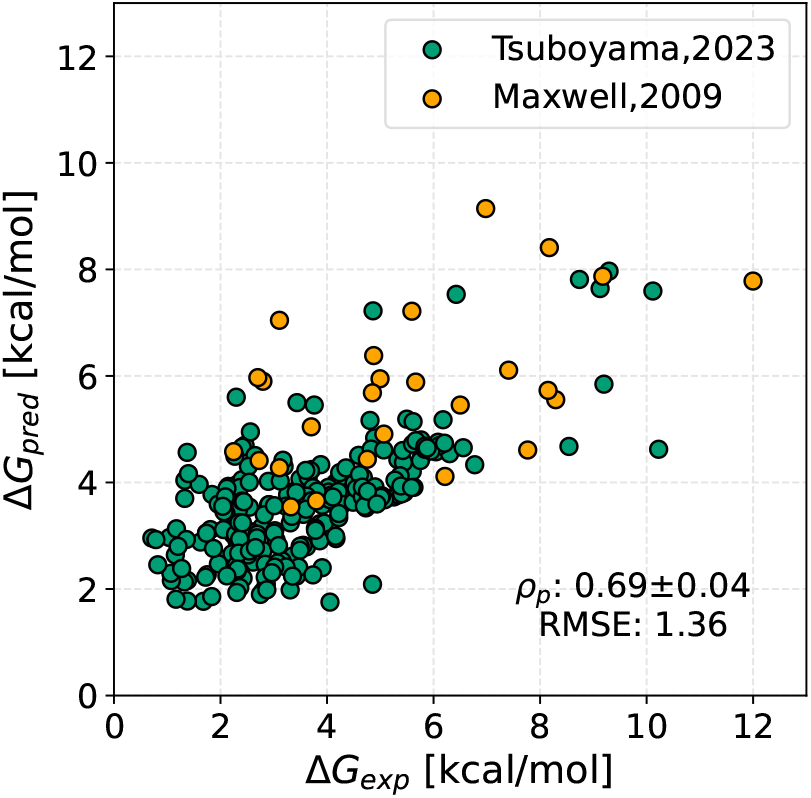
Predicting absolute protein stability using a generative model. We show a comparison between predicted and experimental stability values for 265 proteins from two different data sets (represented by different colours). The predicted values are based on Eq. 1 and have been rescaled to units of kcal/mol (Δ*G*_pred_). The Pearson’s correlation coefficient (*p*_*p*_) and the RMSE (in kcal/mol) are shown. The standard error for the correlation coefficient was obtained by bootstrapping.

Having assessed that the summed likelihoods correlate with global stability, we investigated whether absolute stability was correlated with the summed likelihoods of specific amino acid types or groups of residue types. We found that the summed likelihoods of two aliphatic residue types, valine and isoleucine, correlated the strongest with the global stability (Fig. 3A), and the sum of these two contributions correlate strongly with global stability (*ρ*_*P*_ : 0.64, Fig. 3B). When we aggregated the amino acid likelihoods over all the hydrophobic amino acids, the results correlate with the global stability as much as the total sum of the likelihoods (*p*_*P*_ : 0.67, Fig. 3C), in line with the assumption that protein stability is largely determined by the hydrophobic effect; the correlation using only buried residues was slightly lower (*p*_*P*_ : 0.63, Fig. 3D). Together, these results show that the summed model likelihoods correlate with global protein stability, and that this correlation to a large extent is driven by the hydrophobic amino acids. We note that these correlations are all obtained by representing the protein structure solely via the polypeptide backbone.

**Figure 3.**
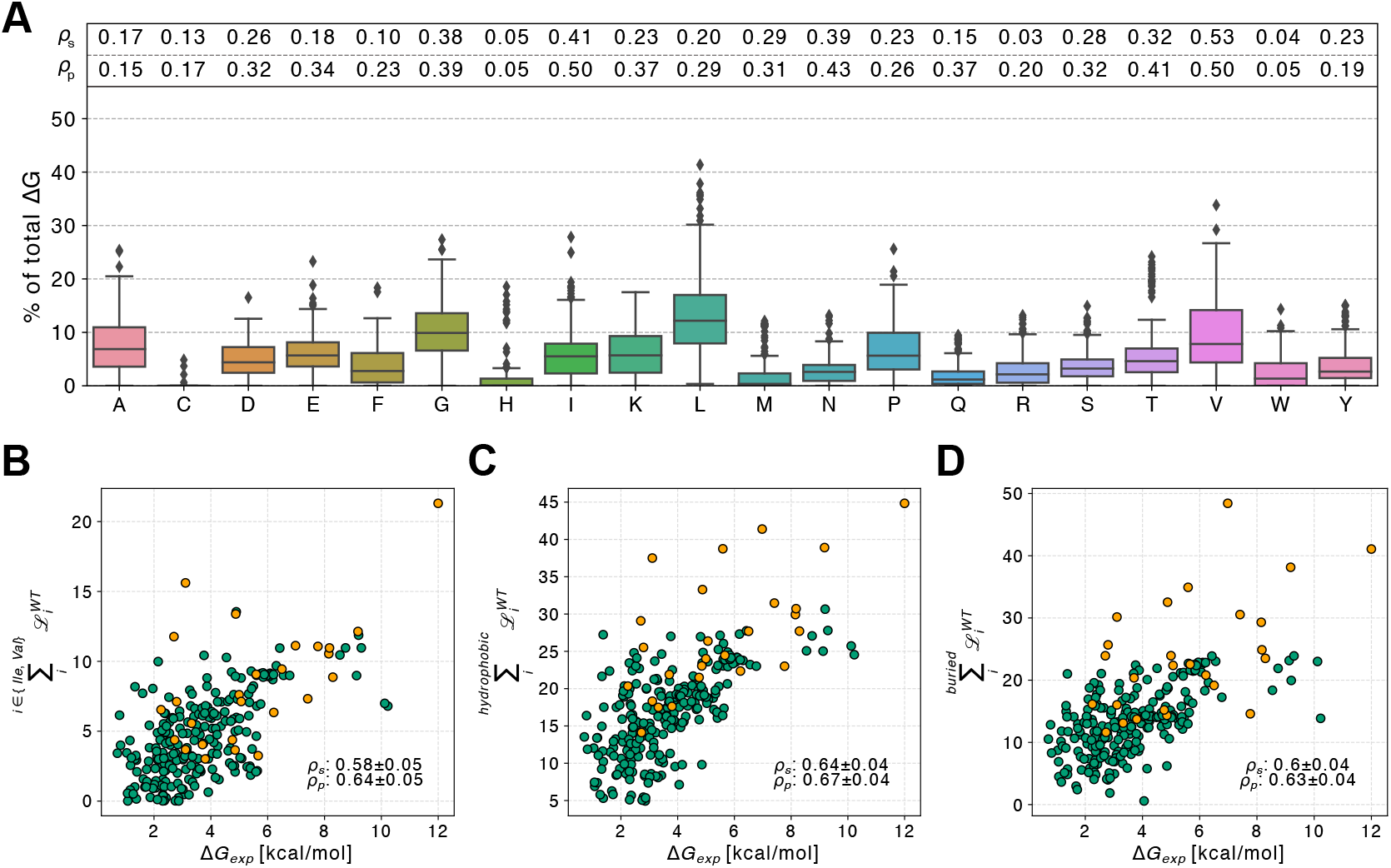
Accuracy of absolute stability predictions using amino acid subgroups. (A) Box plots show the contribution of different amino acid types to the overall absolute stability for each of the 265 proteins in our benchmark set. The Spearman’s and Pearson’s correlation coefficients with the experimental measures are shown at the top of each amino class boxplot. We also show the correlation between the Δ*G*_exp_ values for the 265 proteins in our dataset and the absolute stability calculated from the summed likelihoods of (B) valines and isoleucines only, (C) hydrophobic residues only, and (D) buried residues only. In each panel we show Spearman’s and Pearson’s correlation coefficients and standard errors obtained by bootstrapping.

Having shown that our approach can be used to predict absolute folding stability at a level greater than simpler baseline models, we then asked whether other machine learning models than ESM-IF might achieve similar accuracy using the same approach. To investigate this we used Eq. 1 together with likelihoods obtained from (i) a sequence-based language model (ESM-1b) (***Rives et al., 2021***) to assess the amount of information that could be extracted directly from the sequence using a generic transformer architecture or (ii) likelihoods from a different generative model (Chroma; ***Ingraham et al***. (***2023***)) as a comparison between different design architectures. We also tested energy evaluations using the empirical force field FoldX (***Schymkowitz et al., 2005***), which can be used to predict changes in protein stability. Among the methods tested, we find that the ESM-IF likelihoods achieved the highest correlation with experimental stability measurements (Table 2 and Fig. S5). We note that the sequence-based model, when used together with Eq. 1, achieves a near-zero correlation with the stability measures, highlighting that the stability of a given fold is a difficult task to learn from evolutionary conservation information alone. We also note that while FoldX has been shown to be useful for predicting ΔΔ*G* from both experimental (***Schymkowitz et al., 2005***) and predicted (***Akdel et al., 2022***) structures, it was not developed to predict Δ*G*.

**Table 2.** Pearson’s correlation coefficient between the different alternative model tested including ESM-IF, for the predictions of absolute stability on the experimental dataset.

| Method | ESM-1b | FoldX | Chroma | ESM-IF |
| --- | --- | --- | --- | --- |
| $\rho_P$ | 0.25 | 0.04 | 0.62 | 0.69 |

### Prediction of stability differences between different protein conformations

Having shown that we can use the ESM-IF likelihoods to predict absolute protein stability from a given structure, we then asked whether we could use the same model to predict the difference in stability between two distinct conformations (Fig. 4C). By thermodynamic consistency, this corresponds to predicting the free energy difference between two distinct conformational states, each represented by a single backbone structure.

**Figure 4.**
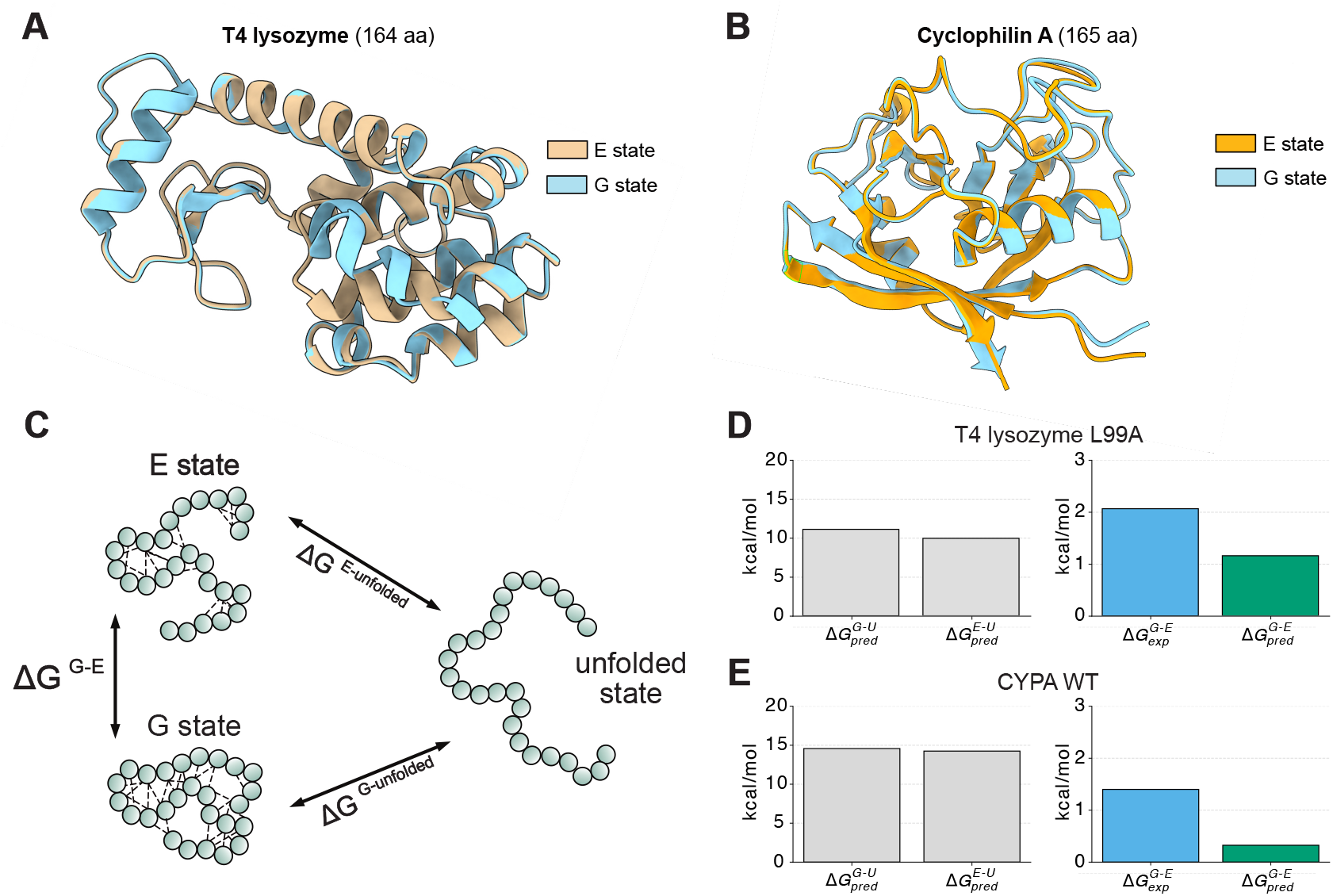
Evaluation of the ability to predict the free energy difference between possible conformations for selected proteins. For proteins with experimentally determined structures of distinct conformational states and the free energy difference between these states, such as the G and E states of (A) T4 lysozyme (T4L) L99A variants and Cyclophilin A (CYPA), we calculate (C) their free energy difference as the difference between their predicted absolute stability. We have predicted and compared with the experimental conformational free energies for (D) T4L and (E) CYPA. In panels D and E, the left plots show the predicted absolute stability for the two states; the right plots shows a comparison of the experimental stability difference and the difference between the predicted absolute folding stabilities.

To assess this ability, we selected two systems where the energetic difference between two states in solution has been measured experimentally using NMR relaxation dispersion techniques. The first system is the L99A variant of T4 lysozyme (T4L L99A, Fig. 4A), which is known to exchange between two conformations, termed G (ground) and E (excited), and with the E state being less stable than the G state by ca. 2 kcal/mol (***Mulder et al., 2001***; ***Bouvignies et al., 2011***). Using structures of the G and E states (***Bouvignies et al., 2011***) as input to ESM-IF and Eq. 1 we predict that the E state is 1.2 kcal/mol less stable than G, in qualitative agreement with experiments (Fig. 4D).

As the second example, we selected Cyclophilin A (CYPA, Fig. 4B) where NMR experiments have shown a free energy difference between two states of 1.7 kcal/mol (***Eisenmesser et al., 2005***), and where models of the two states have been obtained by crystallography (***Fraser et al., 2009***). Again, we used ESM-IF to predict the free energy difference between the two configurations. As for T4L L99A, we also predict a free energy difference whose sign is correct, but with a magnitude (0.3 kcal/mol) that is substantially smaller than found in experiments (1.7 kcal/mol; Fig. 4E). We here note that the ESM-IF model only uses the backbone structure as input, and that it is still not fully known whether the free energy difference in CYPA measured by NMR is fully reflected in the small structural difference (C_*α*_ RMSD of 0.34 Å), between the two structures (***Eisenmesser et al., 2005***; ***Fraser et al., 2009***; ***Papaleo et al., 2014***; ***Wapeesittipan et al., 2019***; ***Thompson et al., 2019***).

While these results demonstrate that our approach cannot yet accurately predict free energy differences between states, they suggest that as prediction accuracy of the stability predictions increase, it might be possible to resolve conformational free energies of the magnitudes found in these two test systems.

### Prediction of the absolute stability of more complex systems

Having shown that ESM-IF provides relatively accurate predictions of absolute stability for small– medium sized proteins that fold in a two-state manner, we asked whether the model could also be used to estimate stability for more complex systems. Specifically, we were interested in assessing whether our approach could also be used to predict the stability of larger, multi-domain proteins—potentially with cooperativity between the different domains—and for proteins where global unfolding does not follow an equilibrium two-state mechanism.

To assess this, we first study two different protein systems, each involving repeated domains: the G5^1^-E-G5^2^ repeat of the Surface protein G (SasG, Fig. 5A; ***Gruszka et al***. (***2015***)) and the Notch ankyrin domain (Nank, Fig. 5B; ***Mello and Barrick*** (***2004***)). In both cases, experimental measures of absolute stability are available for different combinations of the domains (***Gruszka et al., 2015***; ***Mello and Barrick, 2004***). While our approach to predict stability captures the size-dependency across small proteins, we wondered whether it might overestimate stability for larger proteins with (semi-)independent domains. We therefore evaluated the stability for the different domain constructs using either (i) the full structure as input to ESM-IF or (ii) using the sum of stability predictions for the individual domains.

**Figure 5.**
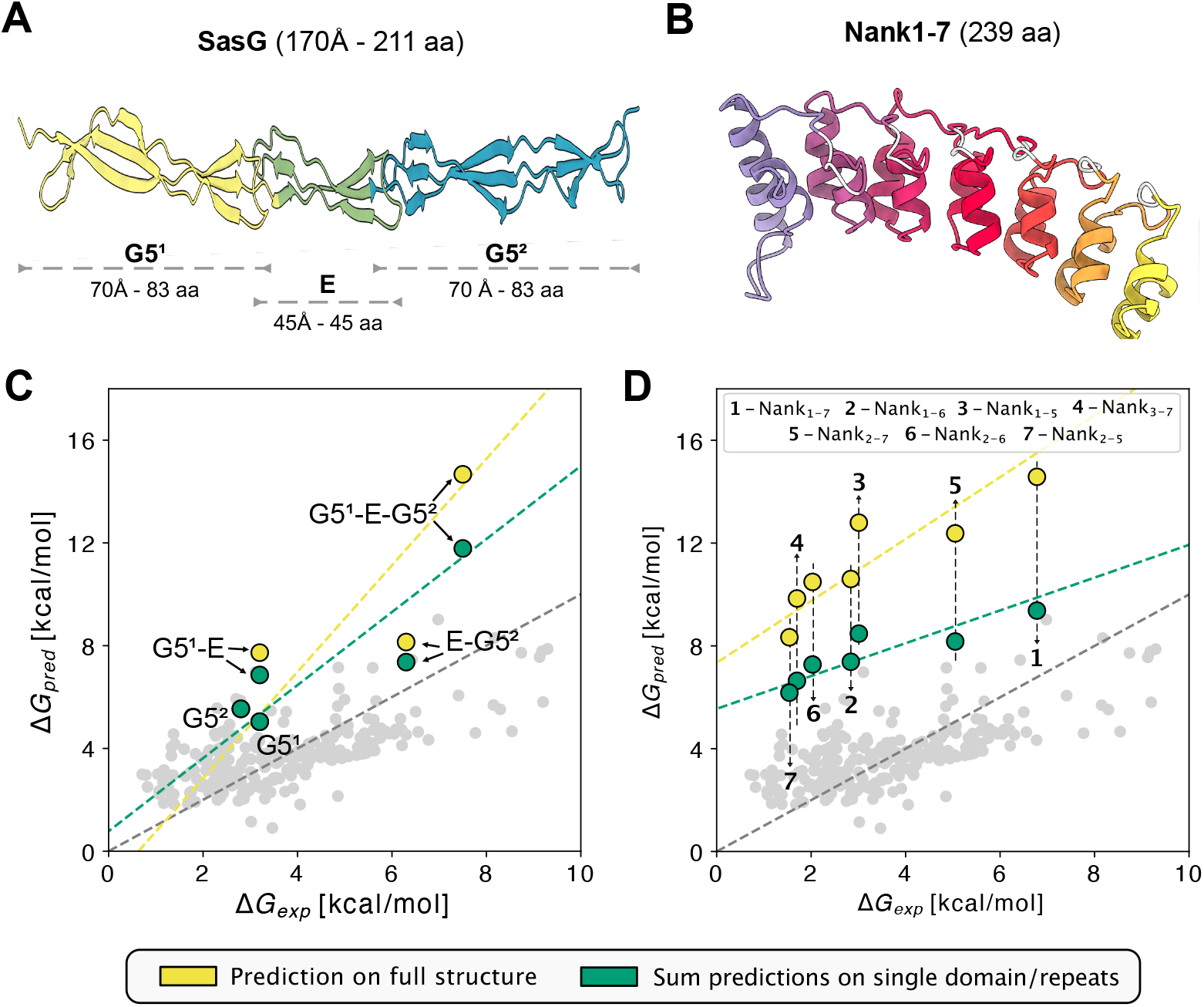
Predicting stability of multi-domain repeat proteins. Starting from (A) SasG and (B) Nank structures we calculated (C,D) the stability of different domain constructs using either structural models with all domains in the construct (yellow) or from the sum of the predictions using the individual domains as input (green). The dashed coloured lines represent the best fit for the specific class of predictions.

For the SasG system we find that the absolute stabilities predicted for the different combinations of domains (G5^1^-E, E-G5^2^, and G5^1^-E-G5^2^) are substantially overestimated when we use the full structure as input to ESM-IF (Fig. 5C). When we instead calculate the stability as the sum of those of the individual domains we obtain results that are closer to experiments, though still overestimated relative to experiments (Fig. 5D). The imperfect agreement may in part originate from the complexity of the system; the E-domain located in between the G5^1^ and G5^2^ domains is disordered on its own and only folds when located between the folded G-domains (***Gruszka et al., 2012***). Thus, the calculations for the G5^1^-E and E-G5^2^ constructs (using models with folded E domains) overestimate the stability. Also, the stability of the G5^1^-E-G5^2^ construct contains a substantial contribution from the interfaces between the different domains (***Gruszka et al., 2015***) that is not captured by our simplified domain-additive approach.

We repeated the analysis for the different combinations of Nank repeats, again predicting the stability both from full structures and as the sum of the stability of the individual repeats. Like for SasG, we find that predictions using the full structures overestimate the absolute stability of the system, whereas stability predictions from summing the stability of the individual repeats were in better agreement with experiments (Fig. 4B). Nevertheless, while the domain-additive model gives results that are in overall better agreement with the magnitude of Δ*G*_exp_, the increase in stability as additional domains are added is not captured accurately. Again, this might reflect that this model does not take into account important contributions from domain interfaces.

Finally, we tested the accuracy of our prediction against proteins with a folding mechanism more complex than two-state folding. We searched the ProTherm database (***Nikam et al., 2021***) for proteins with a multi-state folding process, measured with reversible denaturation and at standard experimental conditions (25 °C, pH 7.0). We found only few systems (less than 10) that met these criteria and selected the two monomeric systems for our analysis: creatine kinase (***Liu et al., 2009***) and exo-small *β* lactamase (***Santos et al., 2007***). Like for the systems described above, our model overestimates the global stability of the two proteins (Fig. S6).

## Conclusions

Protein stability is a key property that underlies protein function, evolution, and our ability to engineer proteins. For example, while accurate prediction of ΔΔ*G* (for one or a few substitutions) might be sufficient when optimizing the stability of a protein, accurate predictions of Δ*G* would enable comparing the stability of a series of orthologues or even completely different proteins. In some sense, prediction of Δ*G* corresponds to predicting ΔΔ*G* between wildly different sequences and structures.

While the prediction of the change in stability due to one or a few substitutions (ΔΔ*G*) is a well-developed area of computational protein science, the prediction of the absolute stability of a protein fold has been a more difficult problem to tackle. One reason is that the global stability contains contributions from a large number of interactions by the protein and solvent in both the folded and unfolded state, making this a difficult quantity to calculate or learn. Also, since ΔΔ*G* is less likely to depend on experimental conditions than Δ*G*, training and benchmarking prediction methods for ΔΔ*G* can use a wider set of experimental data that are collected under different conditions.

Given the relative scarcity of experimental Δ*G* values for distinct proteins under similar conditions, we have here taken the approach of using a pre-trained deep-learning model as basis for our predictions, rather than training or parameterizing a novel biophysical or machine learning model. Nevertheless, we show that there is enough data for benchmarking methods for prediction of absolute stability at physiologically relevant conditions. Here we use the pre-trained model in a relatively simple way (Eq. 1), and while the results are encouraging and useful, we recognize the limitations from the use of the general ESM-IF model and the simple form of Eq. 1. For example, the ESM-IF model represents protein structures via the backbone coordinates providing limitations to the kinds of interactions that are explicitly represented. Our results indicate that the stability of natural proteins is to a substantial extent determined by how likely the amino acid sequence is, with a focus on the side chain packing in the protein chemical environment especially for buried regions of the target fold. While it is not surprising that the environment of the side chains in the structure influences stability, the finding that a simple sum of likelihoods can be used to predict stability of the kinds of proteins we have examined suggests that less stable natural proteins have residues that are suboptimal for their structures. We note that these predictions can be made even though the model does not explicitly represent for example backbone strain or dynamics. While some of these properties may have been learned indirectly during training of ESM-IF, it is also likely that the imperfections in model predictions come from limitations in how the model represents protein biophysics. We also note that the ESM-IF model was trained on the structure and sequences of natural proteins, and that our tests also involve natural (and mostly single-domain) proteins. It thus remains to be tested how accuracy extends to a larger region of the protein sequence and structure universe.

Despite these current limitations, our work shows how the amino acid likelihoods from an ‘inverse folding model’ (ESM-IF) can be used to predict, with useful accuracy, the absolute stability for a series of small–medium sized single-domain proteins with equilibrium two-state folding. As point of comparison, recently developed deep-learning methods for the prediction of ΔΔ*G* achieve an accuracy of ca. 0.5–1.0 kcal/mol for small proteins with larger errors for proteins with more than 100 residues (***Blaabjerg et al., 2023***; ***Dieckhaus et al., 2024***). While these numbers are smaller than the 1.5 kcal/mol that we report here for Δ*G*, we note that the spread of Δ*G* in our benchmark is greater than for ΔΔ*G*. Thus, while substantial work remains to be done, our results demonstrate that absolute stability predictions are possible at a level approaching that of ΔΔ*G*, at least for simpler systems. We expect that representations from deep learning models such as ESM-IF or other structure-aware models, ideally also representing amino acid side chains, may be an excellent starting point for building more accurate and precise predictors in the future.

For larger proteins, including those with multiple domain, our results show more variability. In some cases we find that accuracy improves when predicting the stability via decomposing the protein into individual domains. However, when coupling between domains is substantial, such a divide-and-conquer approach may not work well. In some sense, this is analogous to the predictions of effects of multi-mutants as the sum of contributions of the individual variants, which depends on the additivity of the individual effects. While it may be possible to use our approach also to calculate interface energies between domains, such a decomposition would require automated assessment of how to divide the protein into the relevant domains.

Given the differences we observe between the results on small, simple proteins and larger more complex systems, we caution about making extrapolations about accuracy of Δ*G* predictions from results on small, single domain proteins. We suggest that modelling efforts will benefit both from an increase in the number of Δ*G* values that are measured under consistent conditions, and from making such measurements across proteins of different folds and sizes, and possibly for proteins whose equilibrium folding involves more than two states.

Another interesting avenue for future research is to use approaches such as that described here to examine other questions in protein biophysics. One example we have here explored is to predict the relative stability of different configurations of a protein. We have here tested our approach using two experimentally well-characterized systems where structural models exist for two distinct states. While the free energy differences in the two systems are only slightly larger than the average error of the stability predictions, NMR relaxation dispersion methods can be used to measure free energy differences between conformational states as large as ca. 3 kcal/mol (***Mittermaier and Kay, 2006***). Thus, as prediction accuracy increases we envision that such data will be particularly useful to benchmark methods to predict conformational free energies. We again remind that ESM-IF uses only the backbone structure as input, and so may have difficulties in predicting effects that depend also on differences in side chain packing. Also here, models that represent the atomic structure including side chains might instead be used in the future.

In summary, we have described an approach that uses a generative model to predict absolute protein stability under ambient conditions, and benchmarked it across a number of data sets. While inaccuracies remain, we envision that the ability to predict whether a small–medium sized protein is stable, and by how much, will be useful across many areas of protein science. We also suggest that the data we have collected will be useful to benchmark other approaches to predict stability and other related properties.

## Methods

### Benchmark data

We applied a few filters to the data sets to help ensure the coherence on experimental conditions and types of folding. From the curated dataset (***Maxwell et al., 2005***) we collected 30 protein Δ*G* values and then removed proteins that form obligate homodimers. We also removed the protein VlsE, as it has been shown that despite being a two-state folder protein, it does not follow the usual contact order relationship (***Jones and Wittung-Stafshede, 2003***), reducing the total number of proteins to 26. We note for completeness that the predicted stability of VlsE (18 kcal/mol) is substantially greater than experiment (5.7 kcal/mol), suggesting again that our approach is limited for large proteins.

For the proteins from the previously described ‘megascale’ experiment (***Tsuboyama et al., 2023***), which included 478 unique proteins with estimated Δ*G*, we kept only natural proteins, reducing the size of the data set to 239 proteins. Where a protein Δ*G* was measured in multiple replicates, we used the average of the replicates as the final value.

### Generation of the input structure

As ESM-IF was trained using structures predicted using AlphaFold 2 (AF2) (***Jumper et al., 2021***)), we use LocalColabFold v.1.5.0 (AlphaFold 2.3.1) to generate a structural model for each target fold. We use LocalColabFold with MMseqs2 (***Steinegger and Söding, 2017***) and default parameters including three recycles, no relaxation of the predicted structure, and the maximum multiple sequence alignment size to generate an accurate fold. If an experimentally-derived structure is available for a target protein and conformation, we use it as a template for ColabFold to minimise possible structural deviations introduced by AF2. If the prediction by AF2 differed substantially from this template structure, we performed the prediction again while reducing the MSA size, and thus the influence of the co-evolutionary signal, until the predicted structure results similar to the input structure template (with an RMSD less than 0.5 Å). For the calculations T4 lysozyme we used PDB IDs 3DMV and 2LCB (***Bouvignies et al., 2011***) and for Cyclophilin A we based our calculations on PDB IDs 3K0N and 3K0O (***Fraser et al., 2009***).

### Global stability predictions using ESM-IF

We predicted the global stability value from each structure using the Evolutionary Scale Model ‘Inverse Folding’ model (ESM-IF -model: esm_if1_gvp4_t16_142M_UR50) (***Hsu et al., 2022***)). We used the output logits from the language-model decoder to evaluate the likelihoods for the amino acids at a specific position given the sequence position from the N-term to the preceding position. Specifically, we evaluated the likelihood (ℒ _*k*_) for each amino acid *k* by applying the Softmax on the 20 amino acid logits (*x*) at each position:

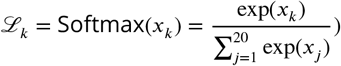

### Other methods for stability predictions

As comparison to the predictions based on ESM-IF we also predicted stabilities using other methods.

#### Absolute contact number

We evaluated the absolute contact number (ACN) of a structure using:

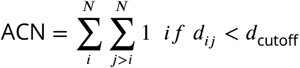

with *N* is the number of residues, *d*_*j*_ is the distance between the *C*_*α*_ atoms of residues and *j*, and *d*_cutoff_ = 8 Å.

#### FoldX

We predicted global stability with FoldX v.5 (***Schymkowitz et al., 2005***) using AF2 predicted structures as input. Specifically, we first used the command ‘RepairPDB’, followed by the command ‘Stability’ to evaluate the different energy contributions and used their sum (total-energy) as absolute stability value. We reverted the sign of this number to represent Δ*G*. We stress here that FoldX was not developed to predict Δ*G*.

#### Chroma

We predicted global stability with the Chroma model (***Ingraham et al., 2023***). Specifically, we used the Github version of Chroma (Dec. 2023) and the design protocol to collect the conditional likelihoods at diffusion time t=0.5. We then use the likelihoods for the wild-type residues and Eq. 1 to evaluate protein stability.

#### ESM-1b

We used ESM-1B (model: esm1b_t33_650M_UR50S) (***Rives et al., 2021***)) to predict absolute stability using only the target protein sequence. For each protein, we collected the output logits of the language model decoder and then used the same approach as for ESM-IF to predict the absolute stability.

### Predictions for Nank and SasG

For both Nank and SasG we first generated AF2 models of the individual repeats using the full structure. For G5^1^-E-G5^2^ we used PDB ID 3TIQ (***Gruszka et al., 2015***) and for Nank used PDB ID 1OT8 (***Mello and Barrick, 2004***). If a loop region was present between the domains/repeats, we included it in the N-terminal fragment. For the additive domain model we simply summed the Δ*G* values predicted for each domain in the construct.

## Data and code availability

Code and data is available at https://github.com/KULL-Centre/_2024_cagiada_stability/

## Acknowledgments

We thank Gabriel Rocklin for comments and suggestions on the manuscript. The research was supported by the PRISM (Protein Interactions and Stability in Medicine and Genomics) centre funded by the Novo Nordisk Foundation (NNF18OC0033950, to K.L.-L.). We acknowledge access to computational resources via a grant from the Carlsberg Foundation (CF21-0392).

## Supplementary Material

### Supplementary Figures

**Figure S1.**
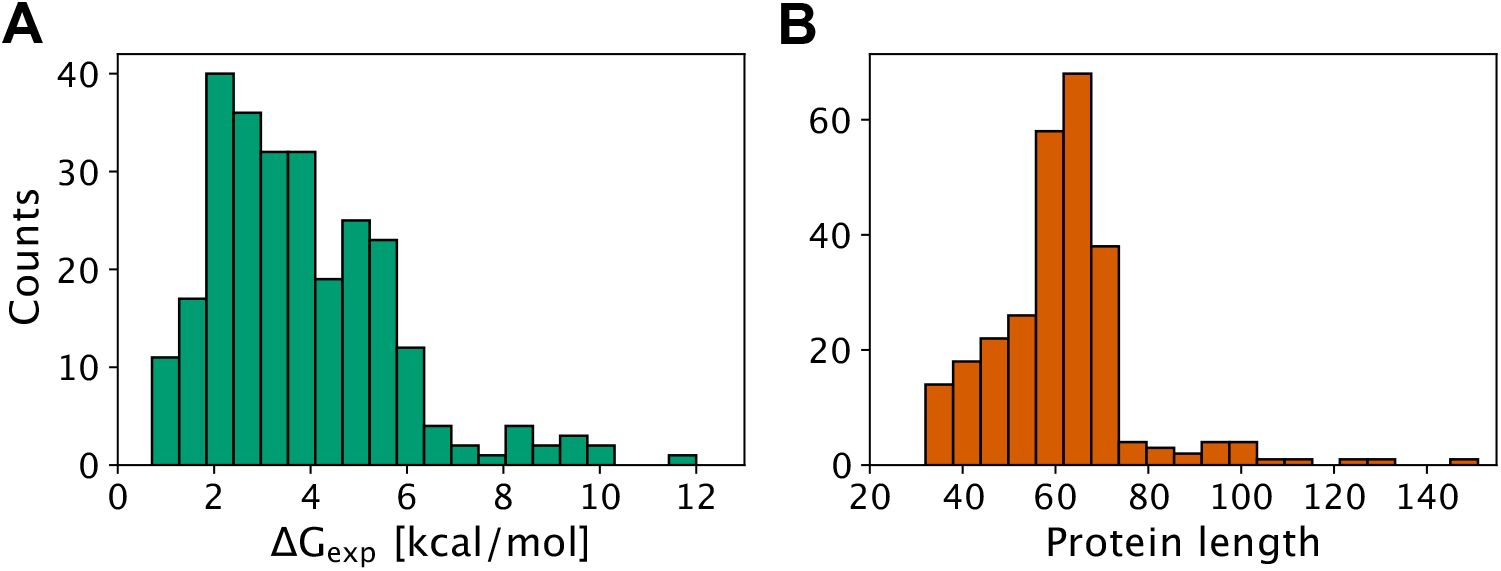
Properties of the benchmark data set. Distribution of (A) Δ*G*_exp_ and (B) number of residues for the 265 proteins in the benchmark data set.

**Figure S2.**
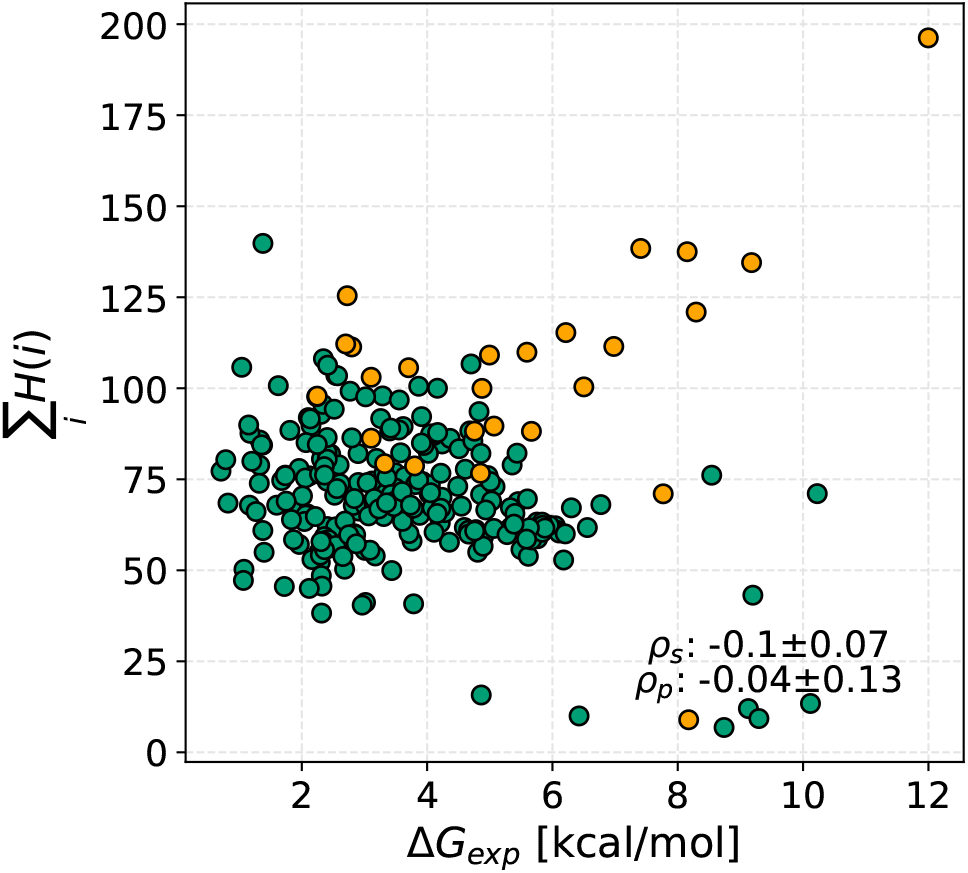
Comparison between absolute protein stability and summed entropy. We show the correlation between the Δ*G*_exp_ (x-axis) values for the 265 proteins in our benchmark dataset and the stability prediction (y-axis) obtained through the sum of positional entropy from ESM-IF amino acid likelihoods. The reported standard error was obtained by bootstrapping.

**Figure S3.**
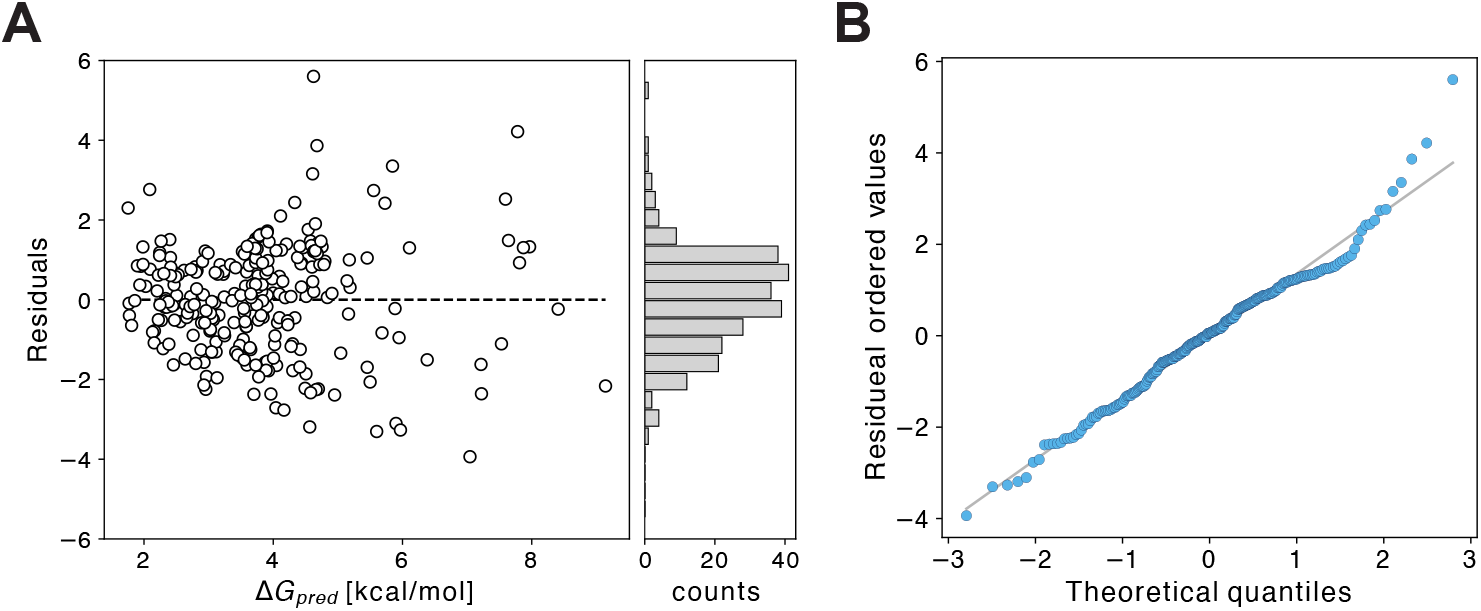
Statistical analysis of the quality of the fit. (A) Residual values for each point during the fit used to convert the ESM-iF score scale into kcal/mol. The residual values are reported on the y-axis for each fitted Δ*G*_pred_. (B) QQPlot (quantile–quantile plot)) for the fitted residuals, suggesting that the errors are roughly normally distributed.

**Figure S4.**
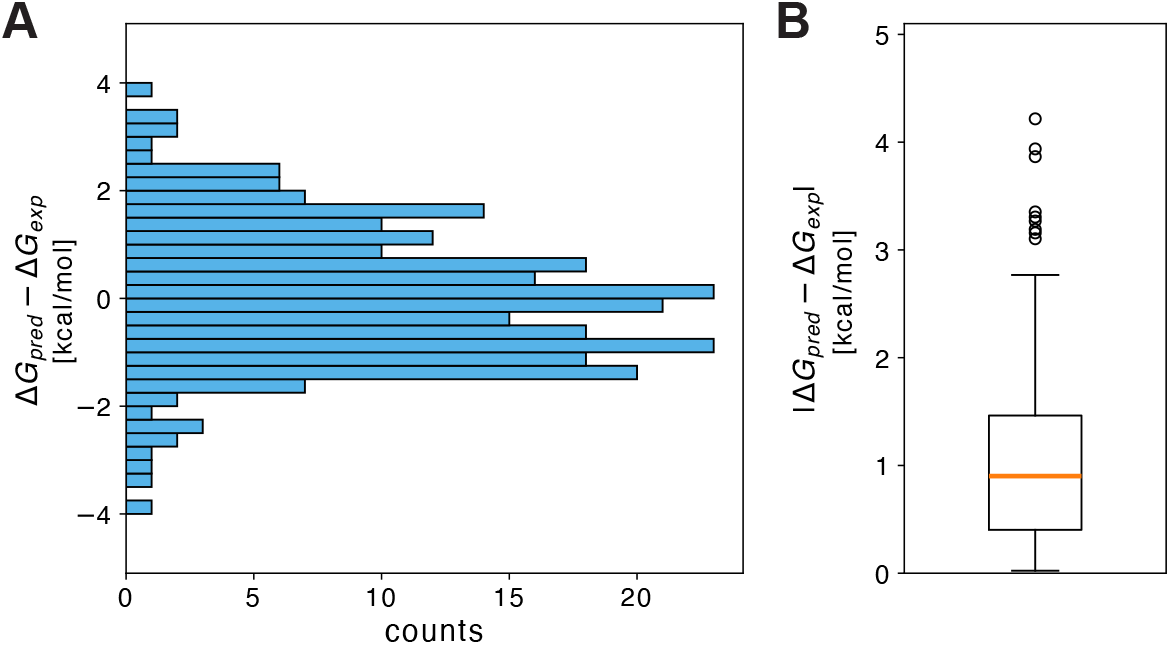
Results of a leave-one-out cross validation of the Δ*G*_*pred*_ predictions. For each of the 265 predictions we fit the conversion from summed likelihoods (Eq. 1) to physical units by fitting on 264 proteins and predicting the Δ*G*_*pred*_ for the left out protein. (A) The distribution of Δ*G*_*pred*_ − Δ*G*_*exp*_ values calculated in this way. (B) Box plot of the absolute difference between the experimental and predicted values. The box plot shows the median as the orange line crossing the box, the lower quartile (Q1, 25%) and the upper quartile (Q3, 75%) as the edges of the box. The lines outside the box represent the one and a half interquantile range (IQR) and the wiskers represent the data points outside the IQR range.

**Figure S5.**
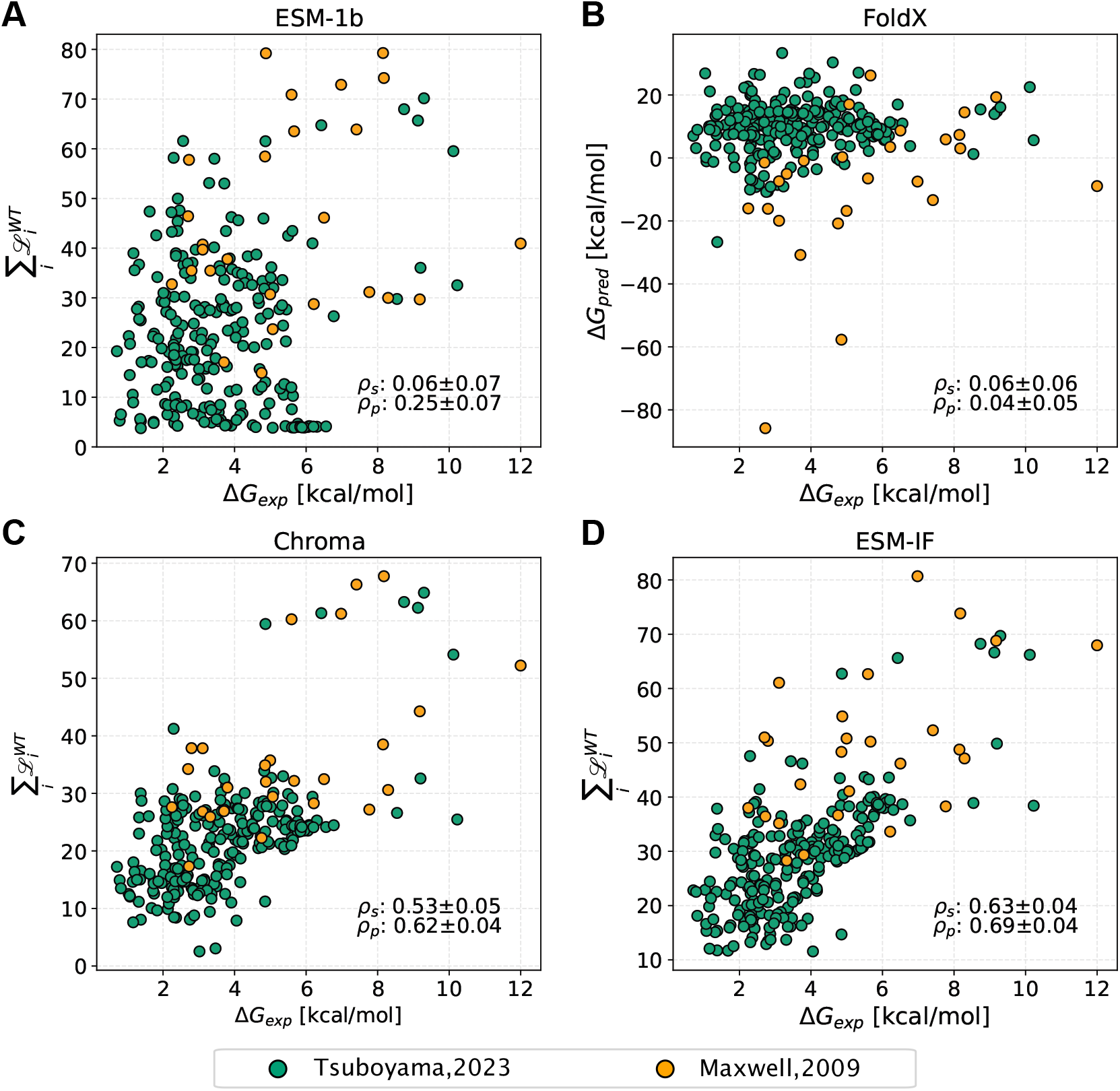
Predicting absolute protein stability using alternative models. We show a comparison between Δ*G*_exp_ values for 265 proteins from two different data sets (represented by different colours). (A) Using likelihoods (*ℒ*) from ESM-1b together with Eq. 1. (B) Total energies from FoldX. (C) Using likelihoods from Chroma together with Eq. 1. For comparison we show (D) Using likelihoods from ESM-IF together with Eq. 1. Panel D is similar to Fig. 1 (without converting to physical units) and is shown here for comparison The Spearman’s and Pearson’s correlation coefficient are shown in each panel (*p*_*s*_, *p*_*p*_). The reported standard errors were obtained by bootstrapping.

**Figure S6.**
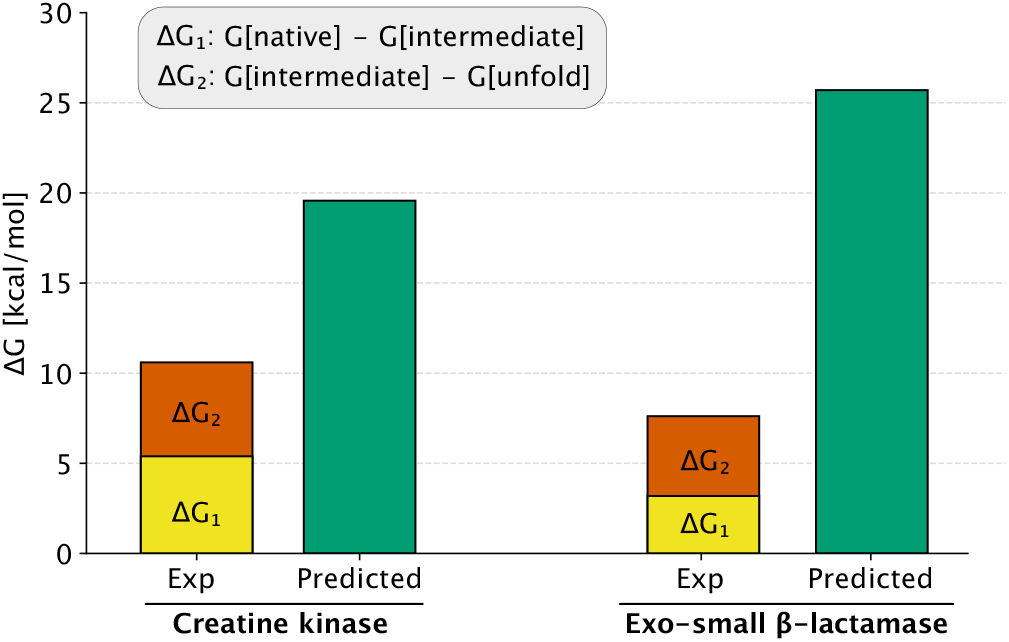
Stability prediction for non-two-state folding proteins. We compared the predicted Δ*G* (green) with the experimental free energy differences going from the native to intermediate states (yellow) and from the intermediate state to the unfolded state (red). Results are shown for (left) creatine kinase and (right) the exo-small *β*-lactamase.

